# Sucrose synthase activity is not required for cellulose biosynthesis in Arabidopsis

**DOI:** 10.1101/2021.02.16.431385

**Authors:** Wei Wang, Sonja Viljamaa, Ondrej Hodek, Thomas Moritz, Totte Niittylä

## Abstract

Biosynthesis of plant cell walls requires UDP-glucose as the substrate for cellulose biosynthesis, and as an intermediate for the synthesis of other matrix polysaccharides. The sucrose cleaving enzyme sucrose synthase (SUS) is thought to have a central role in UDP-glucose biosynthesis, and a long held and much debated hypothesis postulates that SUS is required to supply UDP-glucose to cellulose biosynthesis. To investigate the role of SUS in cellulose biosynthesis of *Arabidopsis thaliana* we characterized mutants in which four, or all six Arabidopsis *SUS* genes were disrupted. These *sus* mutants showed no growth phenotypes, vascular tissue cell wall defects or changes in cellulose content. Moreover, the UDP-glucose content of rosette leaves of the sextuple *sus* mutants was increased by approximately 20% compared to wild type. It can thus be concluded that cellulose biosynthesis is able to employ alternative UDP-glucose biosynthesis pathway(s), and thereby the model of SUS requirement for cellulose biosynthesis in Arabidopsis can be refuted.

## Introduction

Cellulose, the main component of plant cell walls, is synthesized at the plasma membrane by the heteromeric cellulose synthase (cesA) rosette complex, which uses UDP-glucose as a substrate (McFarlane *et al*., 2014). Early evidence from cotton fibres pointed to an important role for sucrose synthase (SUS) in sucrose cleavage and UDP-glucose supply to cellulose biosynthesis. Significant proportion of SUS in the cotton fibres was associated with the membrane fraction, and *in situ* immunolocalisation suggested SUS to be localised in the plasma membrane along the orientation of cellulose microfibrils (Amor *et al*., 1995; Haigler *et al*., 2001). These observations gave rise to a popular model which depicts SUS in association with the cesA complexes channelling UDP-glucose for cellulose biosynthesis (Haigler *et al*., 2001; Stein & Granot, 2019).

Subsequent observations in different plant species supported the association of SUS and cesA complexes. A SUS antibody labelled reconstituted Azuki bean (*Vigna angularis*) cesA complexes in an immunogold labelling assay, and it was reported that this SUS association with the rosette-like structures was required for *in vitro* cellulose biosynthesis activity (Fujii *et al*., 2010). Cotton plants with suppressed SUS activity exhibited reduced initiation and elongation of the cellulose-rich seed fibres, also supporting a SUS function in cellulose biosynthesis (Ruan *et al*., 2003). Immunoprecipitation of cesA complexes from developing wood of *Populus deltoides* x *canadensis* hybrid identified two SUS isoforms further suggesting a direct interaction between cesAs and SUS also in this species (Song *et al*., 2010).

Arabidopsis genome contains six *SUS* genes. Evidence challenging the role of SUS in cellulose biosynthesis was obtained by Barratt *et al*. (2009), who showed that the cellulose content in stems of quadruple Arabidopsis *sus1sus2sus3sus4* (*sus*^*quad*^) mutants did not differ from wild type. However, this conclusion was later questioned by Baroja-Fernandez *et al*. (2012) who maintained that the quadruple Arabidopsis *sus* mutant contained sufficient SUS activity from the remaining two SUS isoforms to support cellulose biosynthesis. Thus, the importance of SUS in providing UDP-glucose to the CesA complexes is yet to be tested unequivocally. To settle the debate about SUS activity in the quadruple *sus* mutant we recently generated lines where all six Arabidopsis *SUS* genes were disrupted (Fünfgeld *et al*., 2021). The Fünfgeld *et al*. (2021) study focused on addressing the role of SUS in ADP-glucose and starch biosynthesis, while here we have used the same *sus* mutants to assess SUS function in UDP-glucose and cellulose biosynthesis.

## Materials and methods

### Plant material and growth conditions

The *Arabidopsis thaliana* ecotype Columbia-0 (Col-0) was used as the wild type control. The quadruple *sus1234* (*sus*^*quad*^) mutant was described in Barratt et al. (2009). The sextuple mutants *sus12345*^*1*^*6* (*sus*^*sext1*^) and *sus12345*^*2*^*6* (*sus*^*sext2*^) were generated by CRISPR/Cas9 as described in Fünfgeld et al. (2021). For the hypocotyl elongation assay, seeds were surface sterilized with 70% ethanol and 0.1% Tween-20 and sowed on ½ MS plates with 1% sucrose and 0.8% plant agar. Then plates were kept at 4 °C for 3 days in the dark followed by light treatment at 22 °C for 6 h to ensure uniform germination. After this the plates were wrapped in aluminum foil and vertically placed at 22 °C for five days. Images of etiolated seedlings were taken by Canon EOS 650D camera and analyzed by ImageJ (http://www.imagej.nih.gov/ij/). For the other experiments plants were grown in soil at 22 °C with a photoperiod of 16 h light and 8 h dark (long day) or 8 h light and 16 h dark (short day) and 65% relative humidity. Valoya NS12 LED tubes were used and the light intensity is 150 µmol m^-2^ s^-1^.

### Anatomy

Sections of the stems from the bottom part of 10-week-old wild type and mutant plants were stained in 0.1% (w/v) Toluidine blue and rinsed with water for three times. The sections were mounted in water and observed under a Zeiss Axioplan 2 microscope equipped with a Zeiss AxioCam HRc camera. Images were processed and analyzed using ZEN 2 blue edition (Zeiss).

### Cellulose analysis

The bottom 5 cm part of stems from 10-week-old plants grown in long day conditions were used for cellulose analysis. The freeze dried samples were ground in liquid nitrogen with mortar and pestle, and alcohol insoluble residues (AIR) were extracted by sequentially heating the samples in 80% and 70% ethanol at 95°C for 30min, followed by treatment with chloroform:methanol (1:1) and washing with 100% acetone. The samples were resuspended in 0,1 M potassium phosphate buffer (pH 7.0) containing 0,01% sodium azide, and starch was removed by digesting the samples twice overnight with α-amylase (Roche, 10102814001, 10U/µl) at +37 °C, 220 rpm. Cellulose content was measured using the Updegraff method (Updegraff, 1969) followed by an anthrone assay (Scott & Melvin, 1953) to quantify the released glucose.

### LC-MS/MS analysis of UDP-glucose

Rosette leaves of 4-week-old wild type and *sus* mutants grown in short days were harvested and homogenized in liquid nitrogen. 10 mg of each sample was placed into a 1.5 ml Eppendorf tube together with 250 μL of ice-cold extraction medium (chloroform/methanol, 3:7) and incubated at −20 °C for 2 h. After this, 10 μL 50 μM UDP-α-D-[UL-^13^C6]Glucose (Omicron Biochemicals Inc, USA) was added to each sample as an internal standard. Samples were then extracted twice with 200 μL of ice-cold water, the aqueous layers were combined and dried in a freeze-dryer. The dried samples were dissolved in 50 µL of 50% methanol and diluted ten-fold prior to the analysis by liquid chromatography-tandem mass spectrometry (LC-MS/MS). The separation of metabolites was achieved by injecting 3 µL of a sample to a HILIC column (iHILIC-(P) Classic, PEEK, 150 × 2.1 mm, 5 µm, HILICON, Umeå, Sweden) and mobile phase composed of (A) 10 mM ammonium acetate in water pH 9.4, and (B) 10 mM ammonium acetate in 90% acetonitrile pH 9.4 at a flow rate of 0.2 mL/min. Ammonium hydroxide was used to adjust pH of the mobile phase and mobile phase was supplemented by 5 µM medronic acid. The gradient elution program was set as follows: 0.0 min (95% B), 15 min (30% B), 18 min (30% B), 19 min (95% B), 27 min (95% B). The LC-MS/MS system consisted of an Agilent 1290 UPLC connected to an Agilent 6490 triple quadrupole (Agilent, CA, USA). Analytes were ionized in electrospray source operated in the negative mode. The source and gas parameters were set as follows: ion spray voltage −3.5 kV, gas temperature 150 °C, drying gas flow 11 L/min, nebulizer pressure 20 psi, sheath gas temperature 350 °C, sheath gas flow 12 L/min, fragmentor 380 V. Multiple reaction monitoring (MRM) transitions of UDP-Glucose and UDP-Glucose-^13^C_6_ were optimized by using flow injection analysis (Table S1). Quantification of UDP-Glucose was conducted based on calibration curve using UDP-Glucose-^13^C_6_ as an internal standard. The calibration for UDP-Glucose covered range from 15 nM to 10 µM with coefficient of determination *R*^2^ = 0.9958.

## Results

To address the role of SUS activity in cellulose biosynthesis in Arabidopsis we made use of the previously generated quadruple and sextuple Arabidopsis *sus* mutants (Barratt *et al*., 2009; Fünfgeld *et al*., 2021). Readers are referred to Fünfgeld *et al*. (2021) for detailed description of the genotypes and the results showing that the two allelic *sus*^*sext*^*-1* and *sus*^*sext*^*-2* mutants contain no measurable SUS activity. Here, wild-type, *sus*^*quad*^ and the *sus*^*sext*^*-1* and *sus*^*sext*^*-2* were grown on soil in 16-hour light period under controlled climate conditions. Under these conditions no visible growth differences were observed between *sus* mutants and wild type (Fig 1a). Compromised secondary cell wall biosynthesis leads to thinner and weaker cell walls, often evident as collapsed xylem vessels (Taylor *et al*., 2000). Therefore, transverse sections of inflorescence stems were prepared to investigate possible cell wall defects in the vascular bundles and the adjacent interfascicular fibres. Light microscopy of the toluidine blue stained cross sections revealed no anatomical defects in the *sus* mutants (Fig 1b). Deficient primary wall cellulose synthesis interferes with cell expansion visible as reduced hypocotyl elongation in etiolated seedlings (Fagard *et al*., 2000). To inspect possible defects in primary cell wall biosynthesis we compared the hypocotyl length in etiolated 3-day-old wild type, *sus*^*quad*^, *sus*^*sext*^*-1* and *sus*^*sext*^*-2* seedlings, but also in these experiments no differences between the *sus* mutants and wild type was observed (Fig 1c and 1d).

**Fig. 1.**
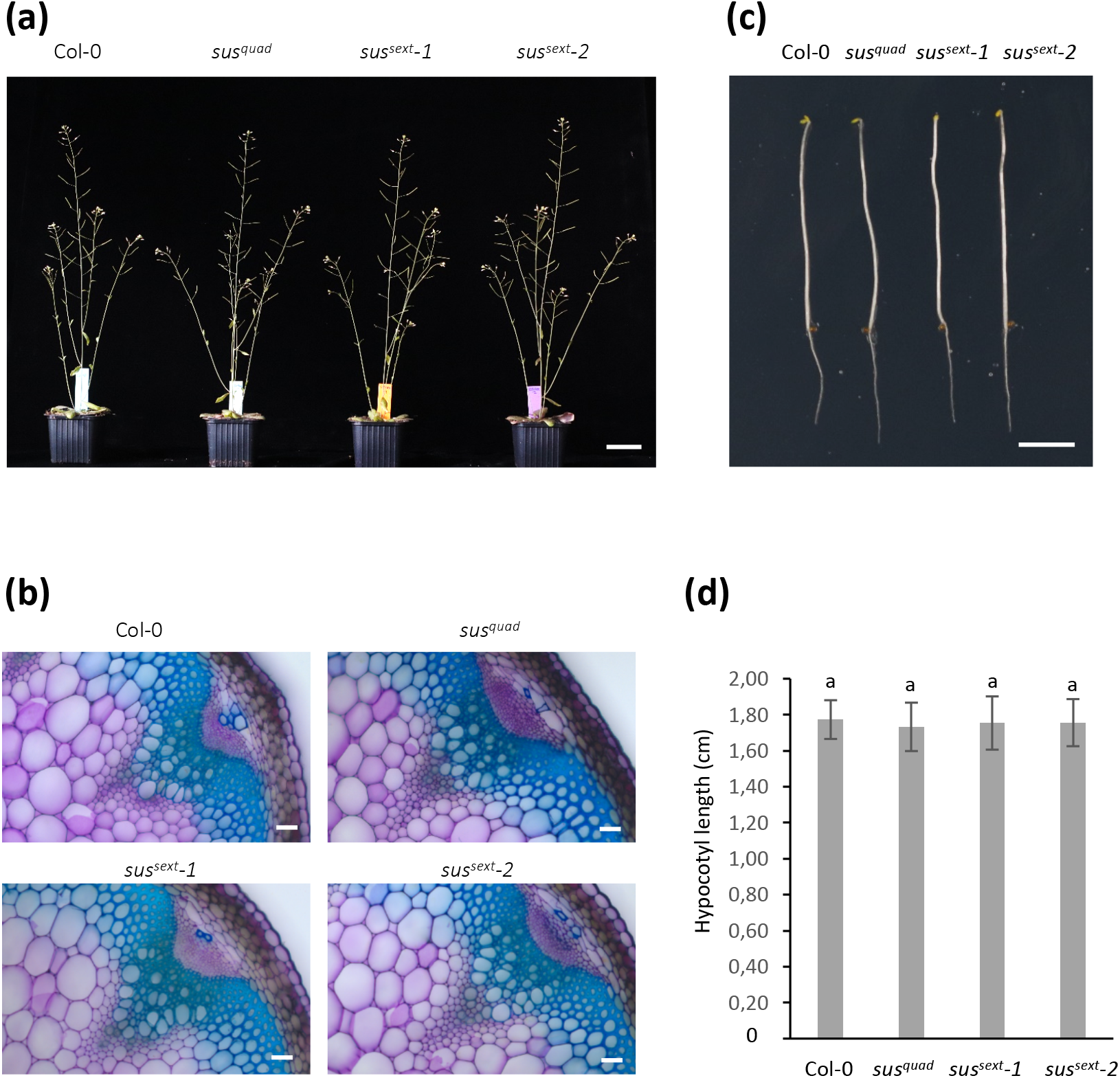
Phenotype of Arabidopsis *sus* mutants. (a) Eight-week-old wild type (Col-0) and *sus*^*quad*^, *sus*^*sext*^*-1* and *sus*^*sext*^*-2* grown in long days (16-h light, 8-h dark). (b) Stem cross sections of Col-0 and *sus*^*quad*^, *sus*^*sext*^*-1* and *sus*^*sext*^*-2* stained with 0.1% toluidine blue. Scale bars 50µm. (c) Etiolated Col-0 and *sus*^*quad*^, *sus*^*sext*^*-1* and *sus*^*sext*^*-2* seedlings grown for five days in the dark. Scale bar 0.5 cm. (d) Hypocotyl length of Col-0 and *sus*^*quad*^, *sus*^*sext*^*-1* and *sus*^*sext*^*-2* seedlings grown for five days in the dark. Values are means ±SE, *n* = 25 biological replicates. One-way ANOVA, post-hoc Tukey HSD test. Letters indicate no significant differences (*P<0*.*05*).

It is possible that lack of SUS activity leads to more subtle effects on cellulose biosynthesis, which would not manifest as anatomical and growth defects. However, quantification of cellulose content in the stems of *sus*^*quad*^, *sus*^*sext*^*-1* and *sus*^*sext*^*-2* revealed no changes (Fig 2a). To further elucidate the reason for the lack of growth phenotypes and cellulose defects we quantified the UDP-glucose pool in rosette leaves of wild type, *sus*^*quad*^, *sus*^*sext*^*-1* and *sus*^*sext*^*-2*. Reliable quantification of UDP-glucose from plant extracts requires rapid quenching of metabolism, extraction and precise LC-MS/MS analysis. UDP-glucose analysis is difficult to perform with success on traditional reversed phase chromatography due to the polarity of the compound. Here we developed a new method combining HILIC-chromatography with addition of medronic acid in the mobile phase, which was previously proposed to improve peak shapes of phosphate-containing compounds (Hsiao *et al*., 2018). Combination of polymer based HILIC column with medronic acid in the mobile phase resulted in improved peak symmetry (Fig S1). Thus, together with MS/MS including carbon-13 labelled UDP-glucose as an internal standard the method provides an alternative for accurate determination of UDP-glucose levels. This analysis revealed that the size of the UDP-glucose pool in the rosette leaves of *sus*^*quad*^ was in fact increased by 10.6% and by 21.0% and 22.4% in *sus*^*sext*^*-1* and *sus*^*sext*^*-2*, respectively (Fig 2B). These counter intuitive results indicated that alternative UDP-glucose synthesis pathway(s) were upregulated and compensated for the missing SUS activity.

**Fig. 2.**
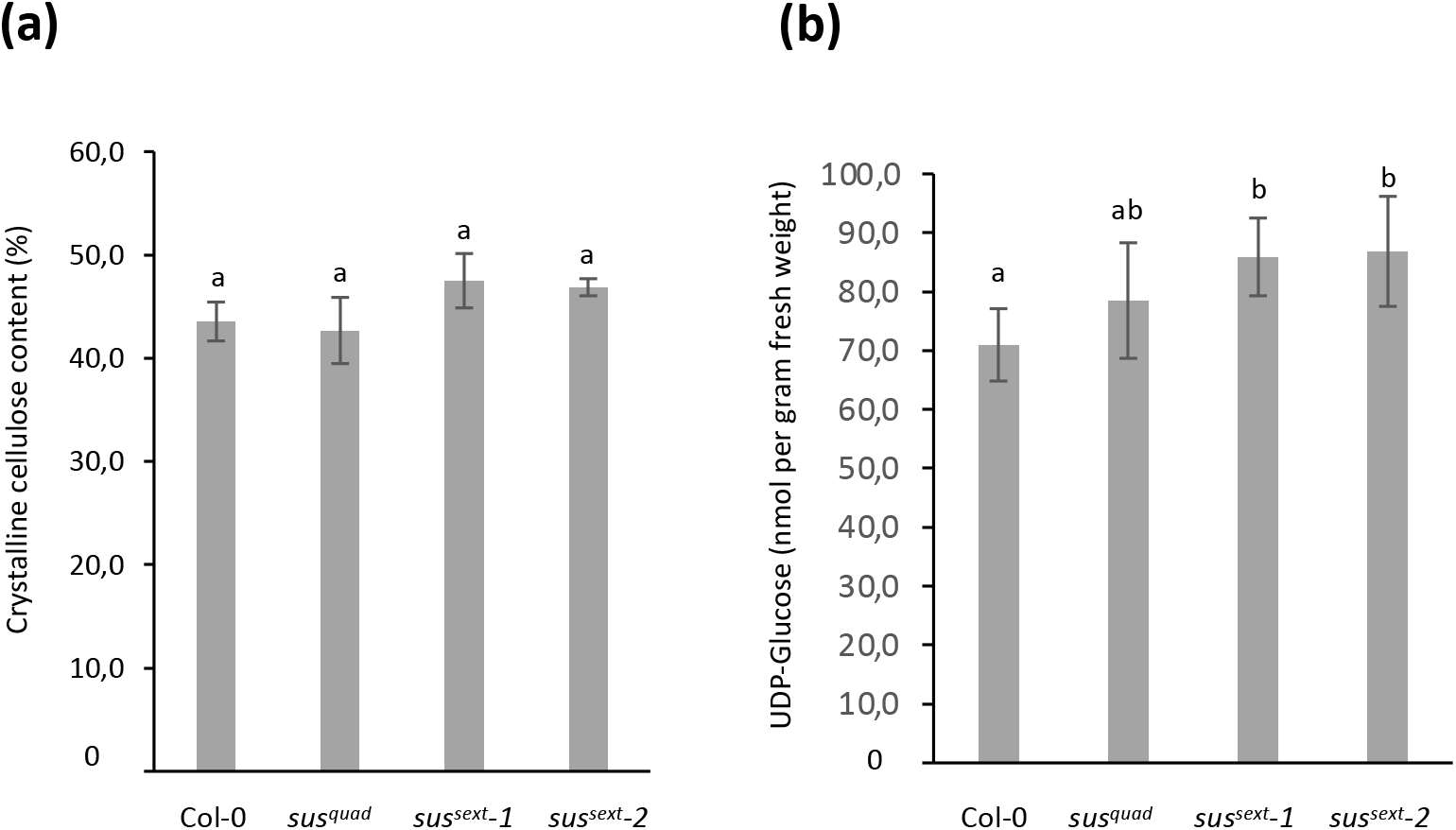
Cellulose and UDP-glucose content in Arabidopsis *sus* mutants. (a) Crystalline (Updegraff) cellulose content in stems of 10-week-old Col-0 and *sus*^*quad*^, *sus*^*sext*^*-1* and *sus*^*sext*^*-2*. Values are means ±SE, *n* = 6 biological replicates. (b) UDP-glucose content in rosette leaves of Col-0 and *sus*^*quad*^, *sus*^*sext*^*-1* and *sus*^*sext*^*-2*. Plants were grown in short days (8-h light, 16-h dark). Values are means ±SD, *n* = 5 biological replicates. One-way ANOVA, post-hoc Tukey HSD test. Different letters indicate significant differences (*P<0*.*05*).

Based on these observations, it can thus be concluded that SUS activity is not required for cellulose biosynthesis in Arabidopsis resolving the long-standing debate about the role of SUS in cellulose biosynthesis. Sucrose cleavage by invertases followed by hexose phosphorylation by hexose-and fructokinases, and UDP-glucose biosynthesis by UGPase activity provide a possible alternative pathway (Barratt *et al*., 2009; Rende *et al*., 2017; Barnes & Anderson, 2018). It should be noted however that Arabidopsis *sus1sus4* mutants exhibit growth defects under hypoxia pointing to the importance of the SUS pathway under stress conditions (Bieniawska *et al*., 2007). A role for SUS in plant stress tolerance was recently also supported by the growth defects observed in hybrid aspen *SUSRNAi* lines grown under natural conditions for five years (Dominguez *et al*., 2021).

## Acknowledgement

We thank Junko Takahashi-Schmidt and the biopolymer analytical facility at UPSC for help with cellulose analysis. This work was supported by Bio4Energy (Swedish Programme for Renewable Energy), the Umeå Plant Science Centre, Berzelii Centre for Forest Biotechnology funded by VINNOVA, and the Swedish Research Council for Sustainable Development (Formas).

## Supplementary material

**Table S1**. LC-MS/MS reaction monitoring parameters.

**Figure S1**. Example of UDP-glucose chromatograms.

