## Supplementary material for "Sucrose synthase activity is not required for cellulose biosynthesis in Arabidopsis"

**Table S1.** Multiple reaction monitoring parameters.

| Analyte | Precursor ion | Product ion | Colision energy [V] | Dwell time [ms] | Type of transition |
|---|---|---|---|---|---|
| UDP-Glc | 565.0 | 322.9 | 25 | 150 | quantifier |
| | | 78.8 | 77 | 150 | qualifier |
| UDP-Glc-$^{13}C_6$ | 571.0 | 322.8 | 21 | 150 | quantifier |
| | | 78.9 | 77 | 150 | qualifier |

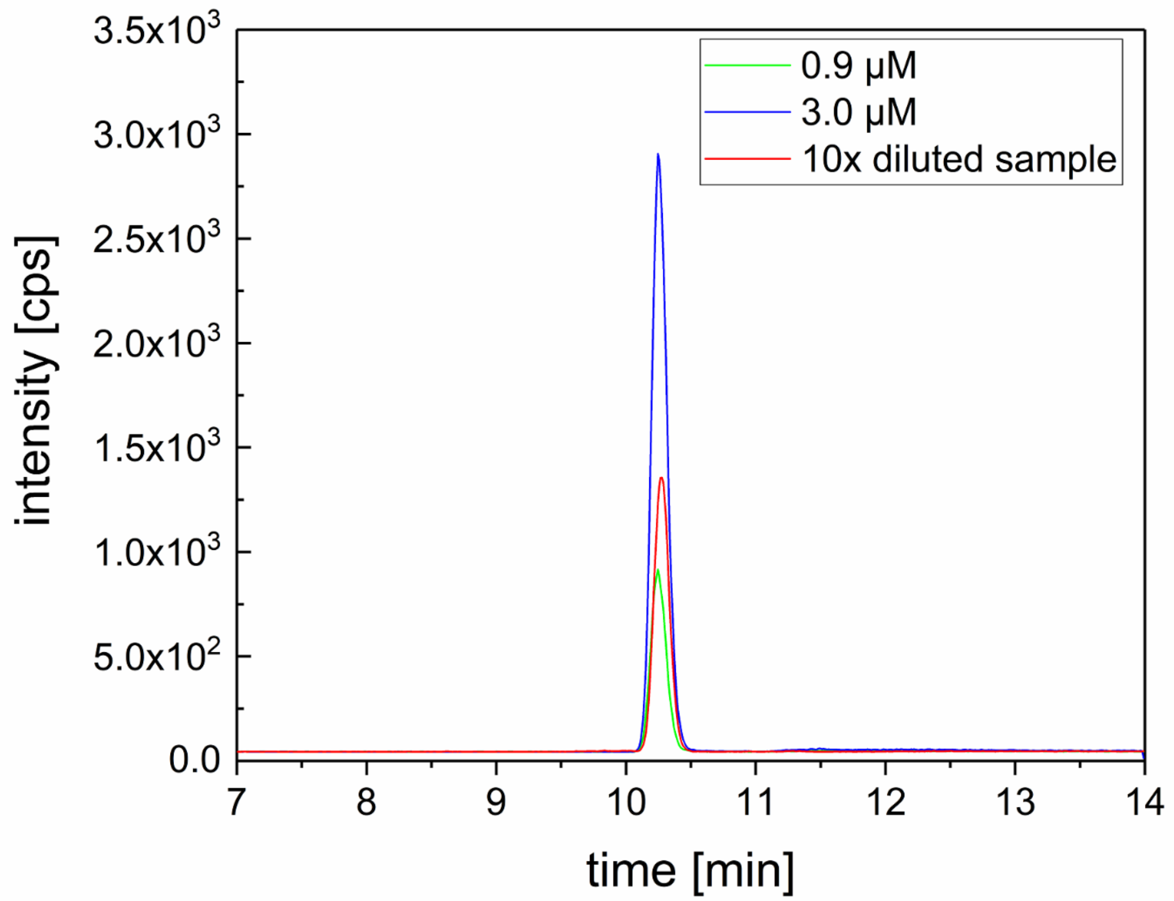


**Figure S1.** Extracted and overlayed chromatograms of UDP-Glucose (MRM transition: 565.0 > 322.9) from calibration solutions and from 10× diluted sample. Chromatographic conditions as described in *LC-MS/MS* analysis section.
